# Multi-Angle Lateral-View Breast Tissue Classification in Thermographic Images Using Deep Reinforcement Learning and Statistical Shape Modelling

**DOI:** 10.64898/2026.10.20.760338

**Authors:** Fatemeh Karimi, Amir Hossein Foruzan, Yen-Wei Chen

## Abstract

This study is a multi-stage method for accurately segmenting and classifying breast tissue in thermographic images, with a specific focus on lateral views captured at multiple angles (30°, 45°, and 90°). Our algorithm is a hybrid boundary detection framework that combines scanning, tracking, and a deep reinforcement learning (DRL) algorithm to delineate the external breast tissue boundary precisely. The DRL agent corrects inaccurately detected landmark points and reduces the average localization error to 17.4 pixels. A Statistical Shape Model (SSM) captures shape variability and refines landmark alignment. A geometric arc-based method estimates the posterior breast boundary, which facilitates robust breast tissue isolation across various lateral view angles. After that, we pull texture features using GLCM (Grey Level Co-occurrence Matrix) and feed them through an SVM classifier. The proposed pipeline boosted diagnostic performance - the SVM hit 74.1% accuracy on our segmented images. It is about 15% better than raw originals and 21% above cropped images. The results prove the effectiveness of our multi-angle lateral view analysis combined with reinforcement learning and statistical shape modelling in enhancing breast cancer detection from thermographic imaging.

## 1. Introduction

Breast cancer is a significant mortality factor for women worldwide. The World Health Organization reports that one in every twelve women may develop breast abnormalities during her lifetime [1]. There are various types of breast cancer with distinct features and treatment considerations. Approximately 70% of breast cancer cases are treatable when the tumor size exceeds 30 millimeters [1]; however, early detection of the malignancy is vital for successful treatment.

Breast Self Examination (BSE) is the simplest method to verify the appearance and feel of the tissue [2]. Mammography imaging is the standard for breast cancer, which takes X-ray images of breast tissue and records density differences in the image. Cancerous tumors exhibit high densities, while the surrounding area shows low densities, making it straightforward to distinguish between the tumor and its surroundings. The primary drawback of mammography is the use of an invasive source that causes genetic changes in the tissue. Other detection techniques are ultrasonography and MRI [3].

Monitoring breast health is equally vital as diagnosis, particularly for early detection of abnormalities. Thermography detects temperature variations in breast tissue using infrared waves. Cancerous tissues typically have higher temperatures due to increased blood flow and metabolic activity. Thermography is a non-invasive screening tool that complements traditional methods, such as mammography [4].

Interpretation of thermography images requires medical specialists who are not likely to be available in rural and underdeveloped regions. Therefore, the need for Computer-Aided Diagnosis (CAD) systems arises. In this research, we accurately classify breast malignancies in thermal images with lateral angles of 30^°^, 45^°^ and 90^°^. This research extend our previous work [5].

## 2. Previous research

### 2.1 Segmentation of thermography images

Segmentation is a basic technique for medical image processing. In thermography images, it improves classification results by focusing on the suspicious areas. There are several methods for segmenting these images.

#### Thresholding and Region-Growing Methods

Díaz-Cortés et al. optimized a multi-level thresholding technique using the Dragonfly algorithm, a metaheuristic inspired by the behavior of dragonflies. This algorithm effectively segments breast thermograms by finding optimal threshold values, thus improving segmentation accuracy [6]. Das et al. focused on accurately segmenting inflammatory and abnormal regions in medical thermal imagery by capturing temperature distributions [7].

#### Edge and Boundary Detection Methods

Mahmoudzadeh et al. introduced a method for detecting boundaries in breast thermograms using a directional version of the Smallest Univalves Segment Assimilating Nucleus (SUSAN) algorithm. It enhances the edges of the breast by incorporating directional information [8]. Additionally, Ensafi et al. precisely separated the left and right breast tissues to distinguish abnormal regions, defining the smallest square as the region of interest [9].

#### Clustering and Superpixel Methods

Sathees et al. combined a variant of fuzzy C-means clustering with anisotropic diffusion filters for detecting abnormalities in breast tissues. Their approach measured the similarity between different regions in breast thermograms to improve the detection of abnormalities [10].

#### Deep Learning Approaches

Sánchez-Cauce et al. utilized a Convolutional Neural Network (CNN) together with clinical data to segment breast thermal images [11]. Tello-Mijares et al. also employed deep neural networks and gradient vector flow for segmenting breast tissue from a frontal viewpoint [12].

### 2.2 Classification Methods for Breast Cancer Detection

#### Machine Learning Approaches

Several researchers have utilized traditional machine learning algorithms to classify breast cancer based on thermal imaging. For instance, Daikushini Satish et al. normalized temperature matrices of thermal images and extracted local energy features using the discrete wavelet transform. They analyzed the differences between the left and right breasts, achieving an accuracy of 91% with a sensitivity of 87.2% using a Gaussian kernel SVM [13]. They also employed first and second-order statistical features, local binary patterns (LBP), and Gabor filters, achieving accuracies of 95%, 92.3%, and 85.4%, respectively. Similarly, Ritam Sharma et al. employed first and second-order statistical features, Gabor features, HOG, and LBP. They used SVM with a radial basis function (RBF) kernel for classification, achieving an accuracy of 96.2% [14].

Moreover, Resmini et al. proposed a hybrid method for screening and detecting breast cancer using dynamic (DIT) and static (SIT) thermographic images. They screened DTI images by the K-Star classifier and achieved an accuracy of 99%, followed by SVM on SIT images, which yielded an accuracy of 95% [15]. Assegie et al. compared SVM and decision tree algorithms for breast cancer prediction, demonstrating that SVM outperformed the decision tree with an accuracy of 93.9% compared to 82.9% [16]. Additionally, Yadlapalli et al. segmented the breast region by thresholding and morphological operations, along with Gabor filters for texture feature extraction, achieving an accuracy of 85.4% in the detection of early breast cancer [17].

#### Deep Learning Approaches

These techniques have significantly advanced the classification of breast cancer. For example, Shobhana Priyasmi et al. distinguished healthy vs. diseased cases by a simple convolutional neural network (CNN) and achieved an accuracy of 77.8% [18]. Mohamed et al. combined the U-Net for segmentation with a custom CNN for classification, resulting in an impressive accuracy of 99.3% in early breast cancer detection [19].

#### Hybrid Models

These models integrate various techniques. Roming Zeng et al. developed the FastLeakyResNet-CIR to enhance the ResNet features and to achieve an accuracy of 98.9% [20]. Additionally, Husaini et al. using the Inception MV4 model, the researchers achieved nearly 100% accuracy by optimizing the number of layers while maintaining computational efficiency [21].

These studies prove the applicability of SVM and deep architectures in breast cancer detection and classification. This review highlights the significant role of breast thermography images, combined with machine learning techniques and image texture analysis, in the early detection of breast cancer. Image normalization and feature reduction techniques, such as Principal Component Analysis (PCA), have further enhanced the performance of diagnostic systems.

In contrast to the predominant attention on frontal view thermography in existing research, our work uses explicitly lateral view images, including angles of 30 degrees and 45 degrees, in addition to the standard 90-degree perspective. Furthermore, we employ a linear kernel in the Support Vector Machine (SVM) for classification.

## 3. The Proposed Method

### 3.1 The outline of our method

Our approach is a multi-stage process to mark and classify the tissue region. Delineation of the breast region involves combining an ad-hoc tracking technique with Statistical Shape Modeling and Reinforcement Learning (RL). This approach enables a robust segmentation of internal and external boundaries of breast tissue across various lateral view angles. Then, we extract appearance features and utilize the SVM to differentiate between malignant and benign lesions.

Key contributions of our methodology include: (1) delineation of the tissue regions in multi-angle lateral views, (2) improvement of classification results by feature extraction from th suspicious region, and (3) robustness of the segmentation technique to image acquisition conditions using the SSM and RL approaches.

### 3.2 Preprocessing

In this study, our algorithm processes both left and right views by flipping left lateral images to obtain the corresponding right lateral view. Then, it transforms the color image into the HSV space. For each image, our method automatically defines ten boundary points along th breast margin. The highest salient point identifies the contact between the chest wall and the armpit, while the lowest salient point is at the beginning of the abdomen.

We select appropriate ranges of Hue, saturation, and value to threshold and identify the breast margins. These masks effectively define the regions of interest; however, they might still contain some noise and unwanted areas. Then, several morphological operations enhance the quality of the masks and eliminate these imperfections.

### 3.3 Boundary Points Detection

In some thermal images, barriers may obstruct the view between the camera and th breast, which can compromise visibility of the breast tissue. Additionally, the patient’s positioning in front of the camera can lead to some regions of the breast being hidden in th captured image. We categorize these instances as incomplete images (Fig. 3a). Conversely, if no obstructions are present, the image is complete (Fig. 3b).

**Figure 1.**
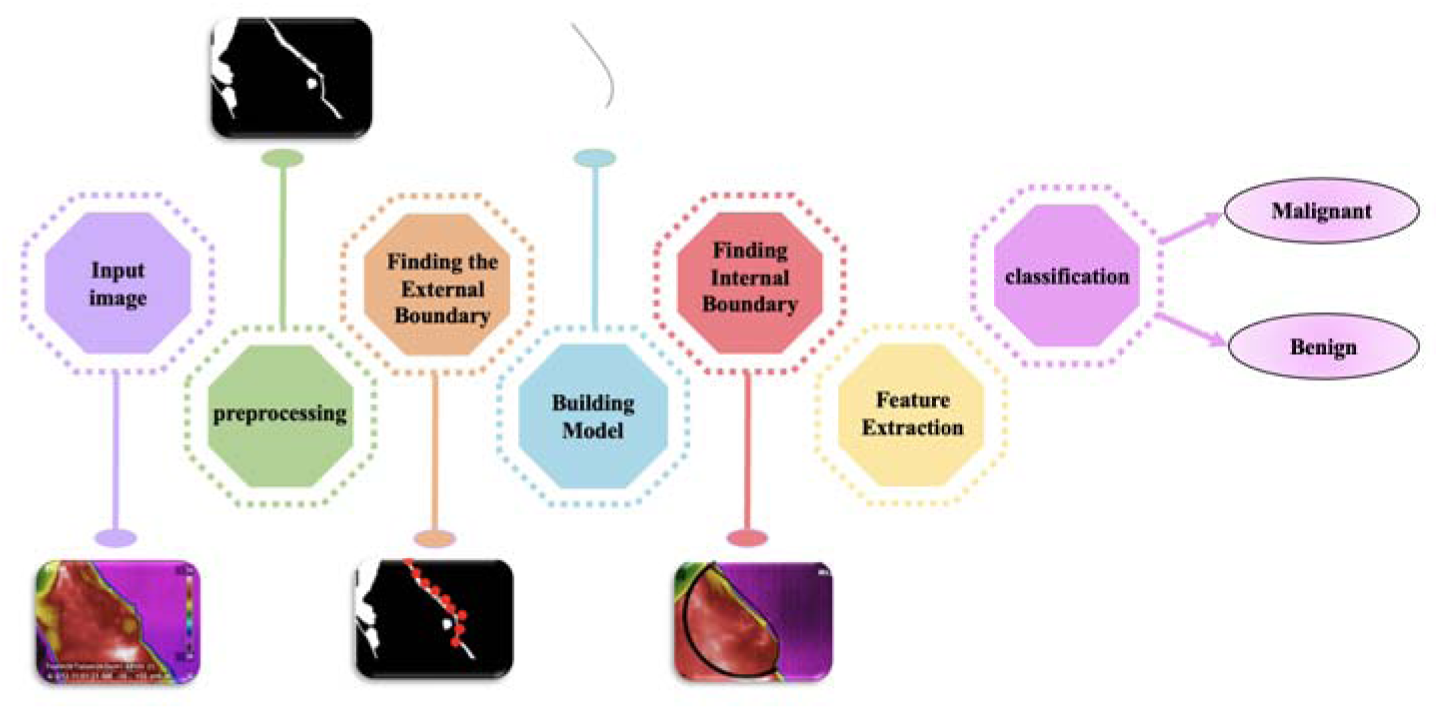
The overview of the proposed method.

**Figure 2.**
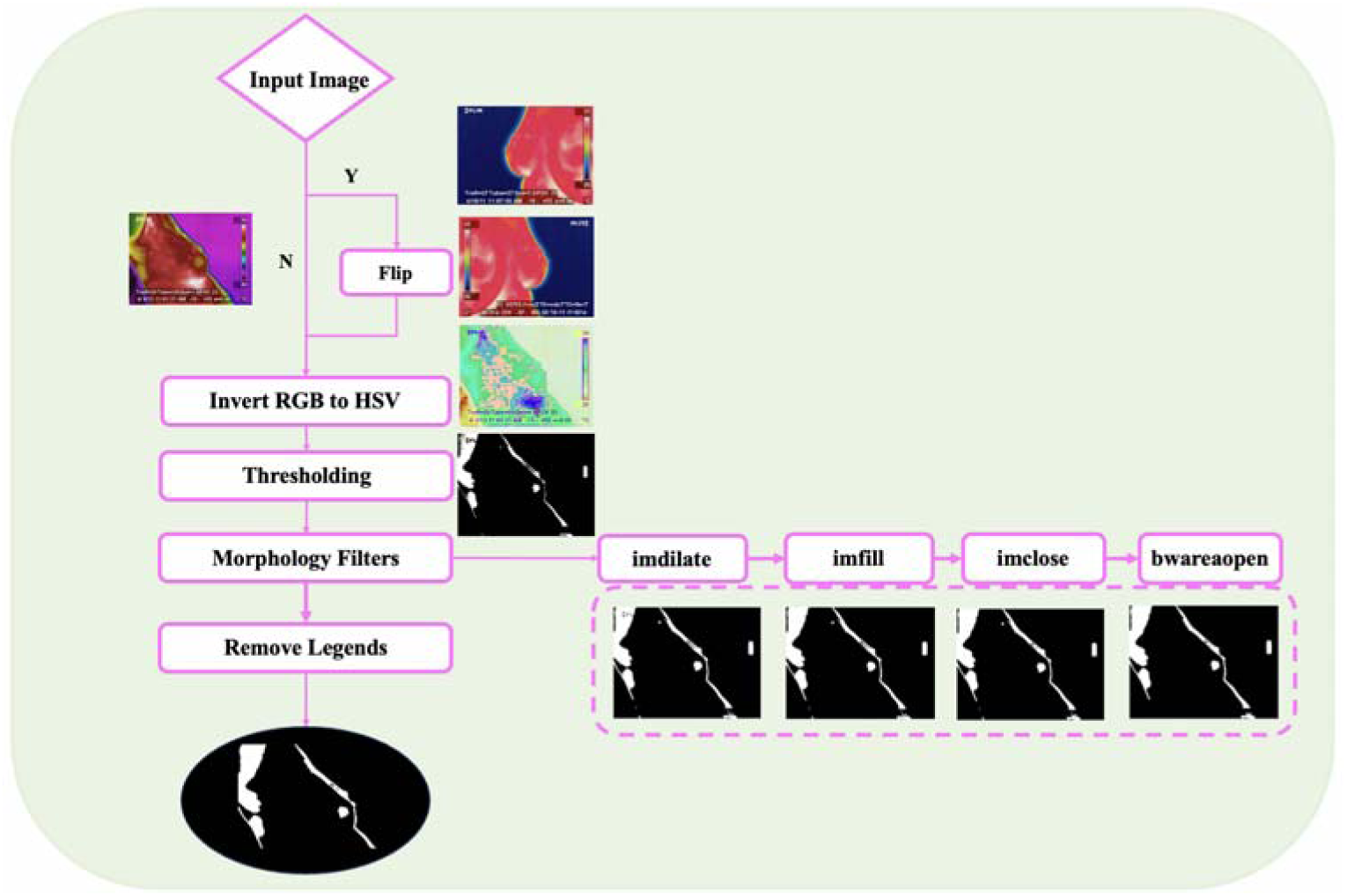
The overview of preprocessing.

**Figure 3.**
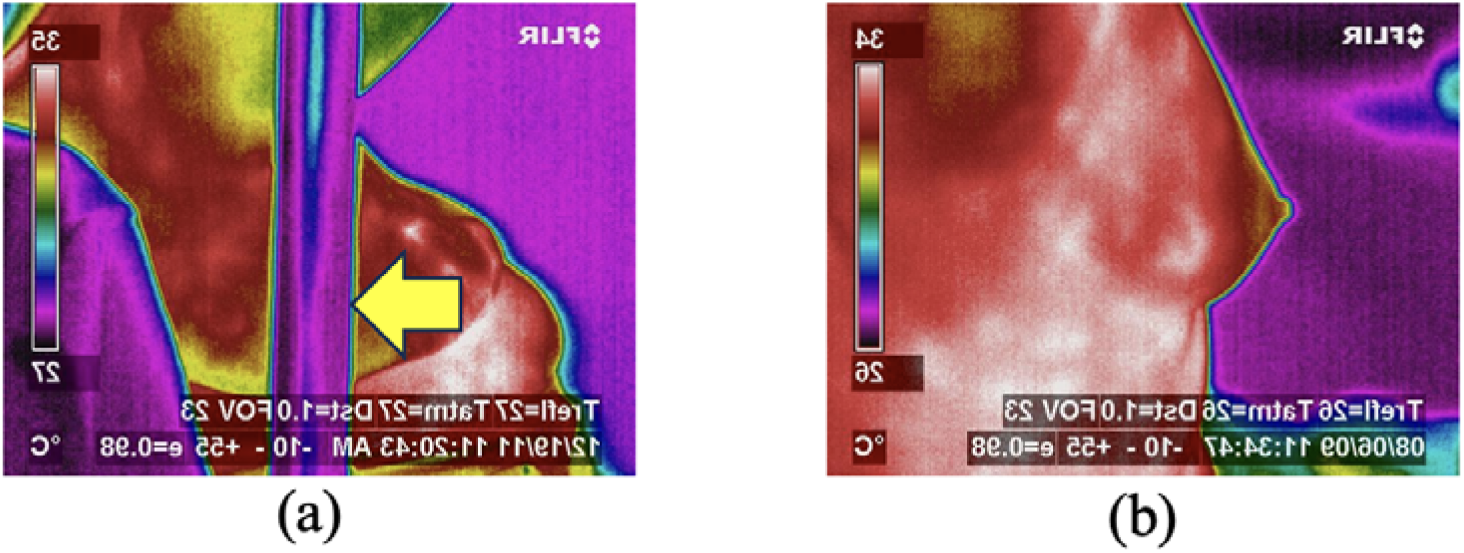
The comparison between (a) incomplete and (b) complete thermal breast images.

The scanning method finds the boundary points starting from the top right corner to the bottom left of the image horizontally and continues to the bottom through a line-by-line search. In case of incomplete images, the result leads to incorrect detection of the boundary points (Fig. 3b). Moreover, the landmarks may be on the arm of the patient or her abdomen. The proposed tracking method improves the results we obtained. The main goal of the tracking approach is to find two salient points of the breast region that are the armpit ( in Fig. 4) and the cross-point of the breast and abdomen ( in Fig. 4). Details of the methods are in [5].

**Figure 4.**
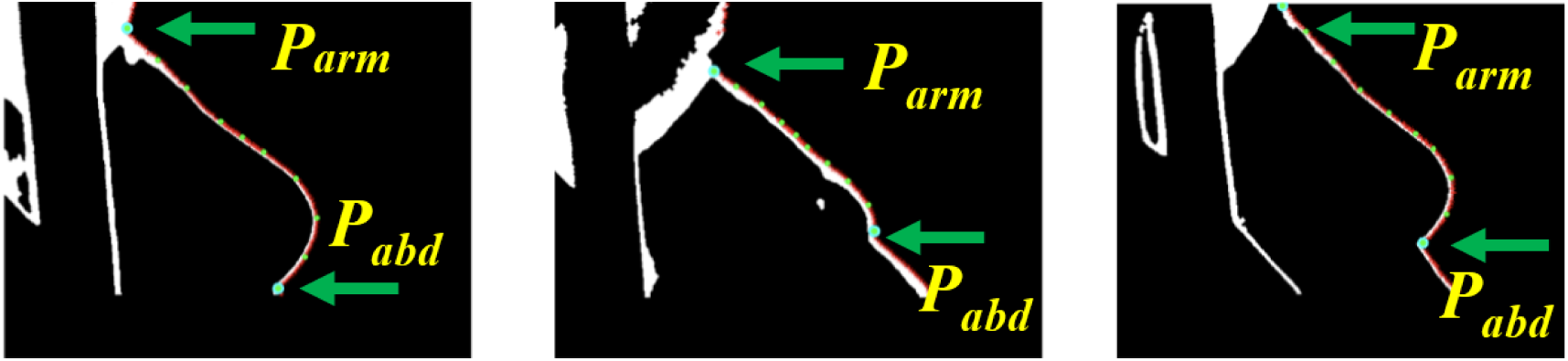
Salient point detection by the tracking method. The tracking method improves the landmark locations obtained by the scanning method.

### 3.4 Boundary Detection by RL Algorithm

We apply reinforcement learning (RL) to improve landmark detection accuracy in challenging cases such as incomplete images. Our RL agent uses a dueling Deep Q-Network (DQN) architecture to make sequential decisions based on local image information. The goal of the RL process is to refine the position of the abdominal salient point, which the tracking algorithm may have inaccurately identified. During training, the agent iteratively minimizes the distance between the erroneously detected point and the correct position.

The DQN processes thermal images of size 400 400 1 pixels, stacked into tensors of shape (10, 400, 400). An initial permutation layer feeds into six convolutional layers using 3×3 kernels and ‘same’ padding for spatial consistency. Filter counts scale progressively from 30 to 120 across these layers, enabling hierarchical feature extraction. We flatten the feature maps and pass them through two separate fully connected streams, value and advantage streams, implementing the dueling architecture. The final Q-values guide the agent’s decisions across four possible actions: up, down, left, and right (Figure 5).

**Figure 5.**
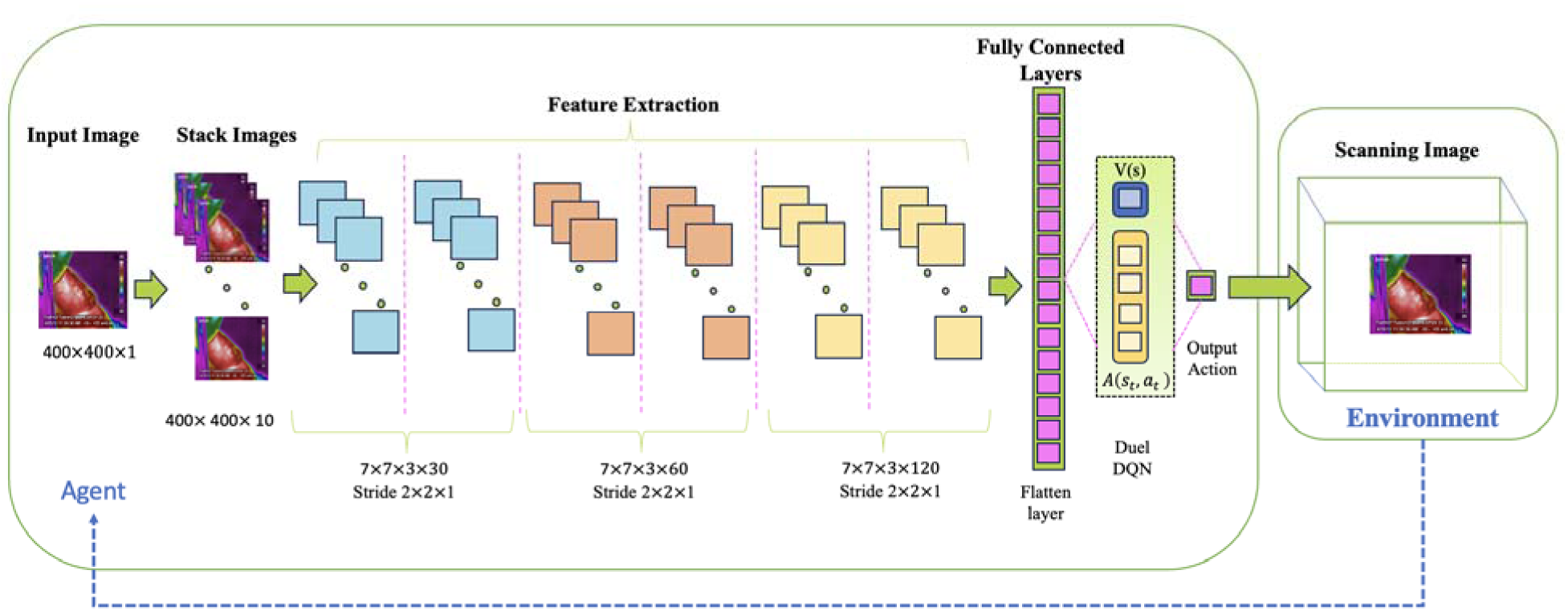
Architecture of the proposed Deep Q-Network (DQN) for salient point correction.

The reward mechanism drives the agent toward the correct landmark position by combining several components:

• **Step Penalty:** -1 to discourage unnecessary moves.

• **Directional Rewards:** positive reward for closing in, penalty for moving away.

• **Proximity-Based Rewards:** Bigger rewards near target; penalties if farther than start.

• **Completion Reward:** A substantial reward is given when hitting the landmark zone.

• **Boundary Penalties:** Cost for leaving the frame.

To enhance the stability and efficiency of the learning process, there are multiple features in the system. In the target network of the architecture, there are two separate networks, one leading network for action selection and one target network for value estimation. The dueling architecture, experience replay, and a balance between exploration and exploitation allow the agent to start with an exploratory policy and gradually transition to a more greedy behavior as it accumulates experience (Figure 6).

**Figure 6.**
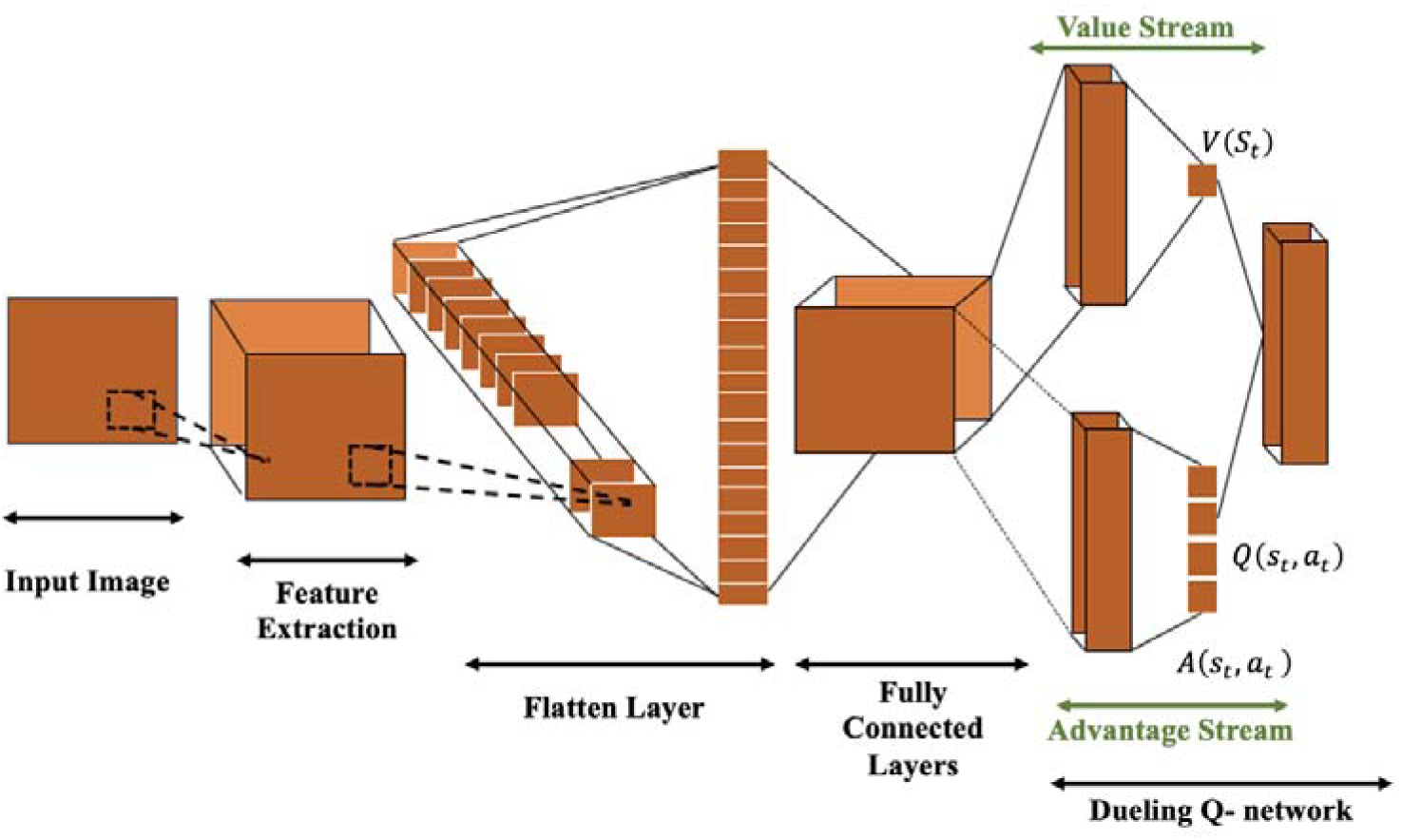
Dueling Q-network structure.

We train our agent with Temporal Difference (TD) learning, where the Bellman equation shapes its cost function (Eq. 1). The learning rate ( ) is set to 0.99, and the discount rate ( ) is 0.95, ensuring the agent considers both immediate and future rewards. The loss function (Eq. 2-3) measures the mean squared error between predicted and target Q-values. Training stability i enhanced using the Adam optimizer with a learning rate of 0.00025 and gradient clipping.

After training completes, our RL agent both refines key points and lays down 10 equally spaced boundary points between landmarks. It handles real-world image variations and landmark shifts effortlessly, proving superior to older methods. By leveraging experience, the RL framework improves landmark detection accuracy and robustness over time.

### 3.5 Building the Statistical Shape Model

Statistical Shape Modelling (SSM) captures the shape variability present in a set of similar objects, such as anatomical structures, by defining a set of landmarks in each training shape. The first step is to remove shape differences resulting from scale, translation, and rotation transforms by the Generalized Procrustes Analysis (GPA). Then, a statistical analysis determine the mean location of each landmark and its variance. Finally, the Principal Component Analysis (PCA) identifies the main modes of variation. Details of the algorithm are in our previous paper [5].

### 3.6 Boundary detection and Feature Extraction of a test image

For a test image, we convert it to gray and then make it binary using the Otsu algorithm. The scanning algorithm defines the initial boundary points, which may be far away from the actual breast border. Then, the SSM algorithm iteratively fits these initial boundary points to the test shape. Fig. 7 shows several cases of breast boundary detection in complete and incomplete shapes.

**Figure. 7.**
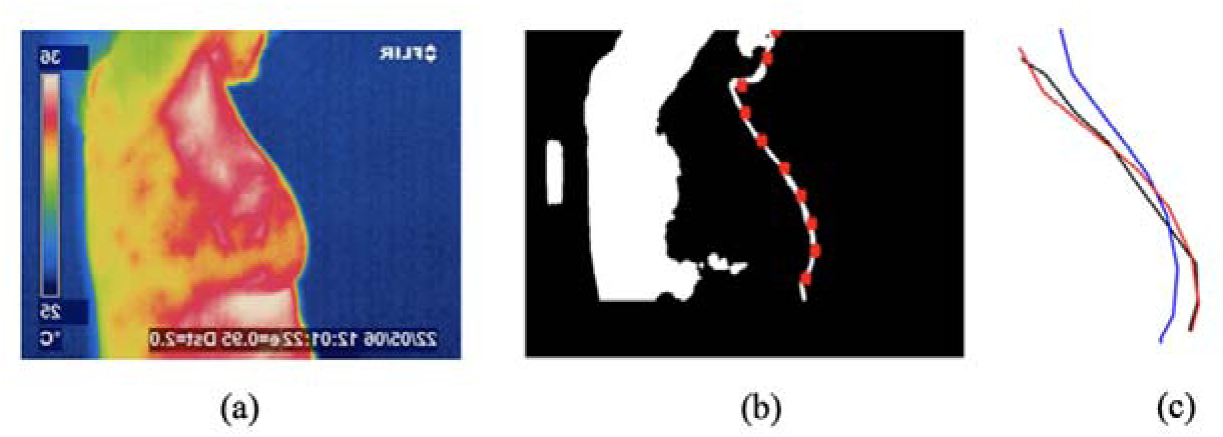
Fitting the SSM model to the landmarks. (a) The row image. (b) The binary image with the landmarks that are shown by red points on the boundary. (c)The black line is the Model that was made by SSM. The blue line is the result of connecting landmark points, and the red line is the fitted model.

We model the internal boundary of breast tissue by a semicircle, which resembles the natural anatomy of breast tissue. Representation of the internal part of the breast by a semicircle ensures that its depth is proportional to the natural size and shape of the tissue. The first and last keypoints described in subsection 3.3 (Parm and Pabd) define the beginning and end of the semicircle, and its center is the mean position of these two points.

After boundary detection of the breast, the Grey Level Co-occurrence Matrix (GLCM) represents the appearance of the tissue. GLCM matrices decode texture patterns, like homogeneity and contrast, in our images. We then feed these features to a linear-kernel SVM, which spots whether tissue is benign or malignant.

## 4. Results

### 4.1 Dataset & implementation Details

The database of our research consists of thermographic breast images acquired at the University Hospital of the Federal University of Pernambuco [22], [23], [24], [25], [26]. These images have undergone clinical classification, falling into one of three categories: cyst, malignant, or benign. The dataset includes both frontal and lateral views of the breast (Fig. 8). Specifically, the lateral views encompass four image types: LEMD (external lateral view of the right breast), LEME (external lateral view of the left breast), LIMD (internal lateral view of the right breast), and LIME (internal lateral view of the left breast). The dataset also has frontal views of the right breast (MD), left breast (ME), and bilateral frontal views with arms down (T1) and arms up (T2). The implementation of the proposed algorithm had two main components. The initial stages, including tracking and Statistical Shape Model (SSM), were developed in a MATLAB environment on a MacBook M2 Pro with 8 GB of dynamic RAM. We customized the SSM using a GitHub repository [27]. The RL implementation used Python 3.6.11, accelerated by parallel processing on two NVIDIA GPUs (RTX 430 and RTX 540). This dual-hardware approach allowed component-specific optimization for computational efficiency.

**Figure 8.**
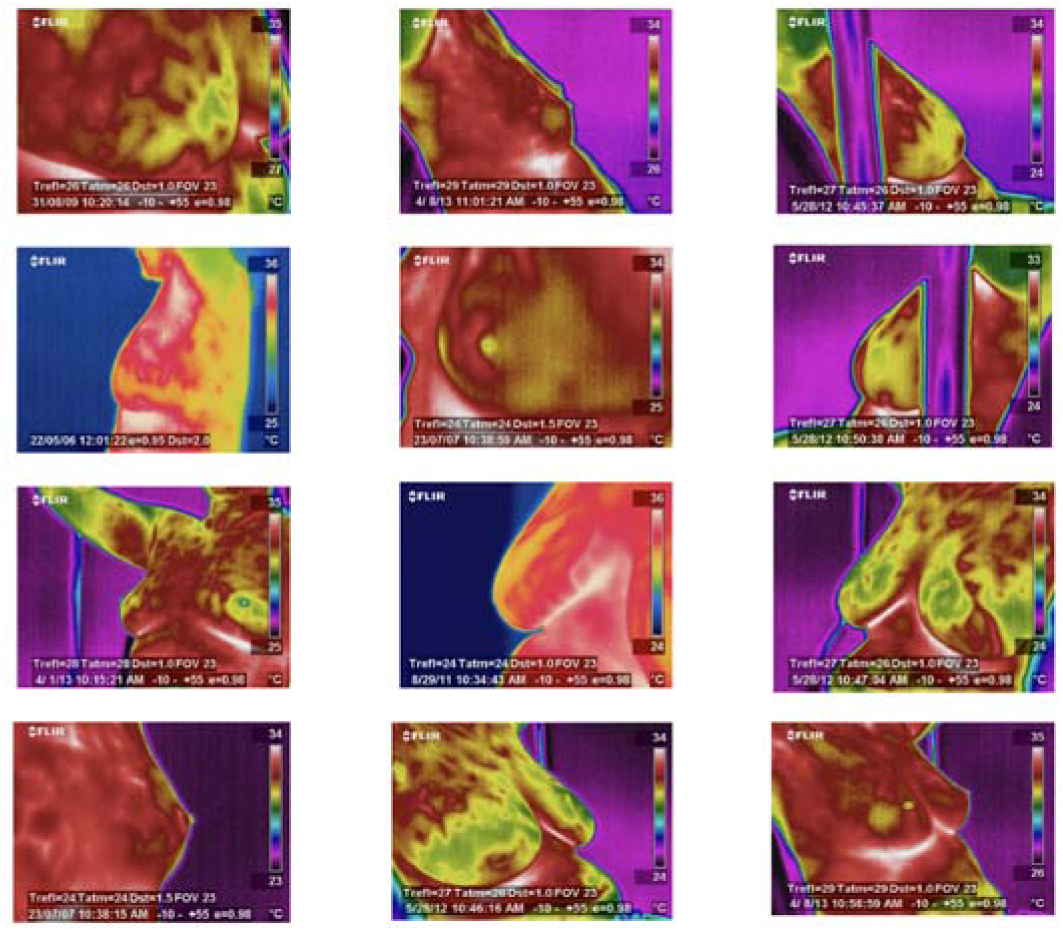
Representative lateral views of thermographic breast images. The data belong to the University Hospital of the Federal University of Pernambuco dataset. These images include LEMD (external lateral of the right breast) in the first row, LEME (external lateral of the left breast) in the second row, LIMD (internal lateral of the right breast) in the third row, an LIME (internal lateral of the left breast) in the fourth row.

### 4.2 Qualitative evaluations

The scanning method frequently locates points on the breast boundary except for *P_arm_* and *P_abd_* (Fig. 4). We developed the tracking algorithm as a remedy; however, the algorithm failed in the presence of obstructions or anatomical variability existed. The RL method addresses these limitations and refines landmark positions further (Fig. 9).

**Figure. 9.**
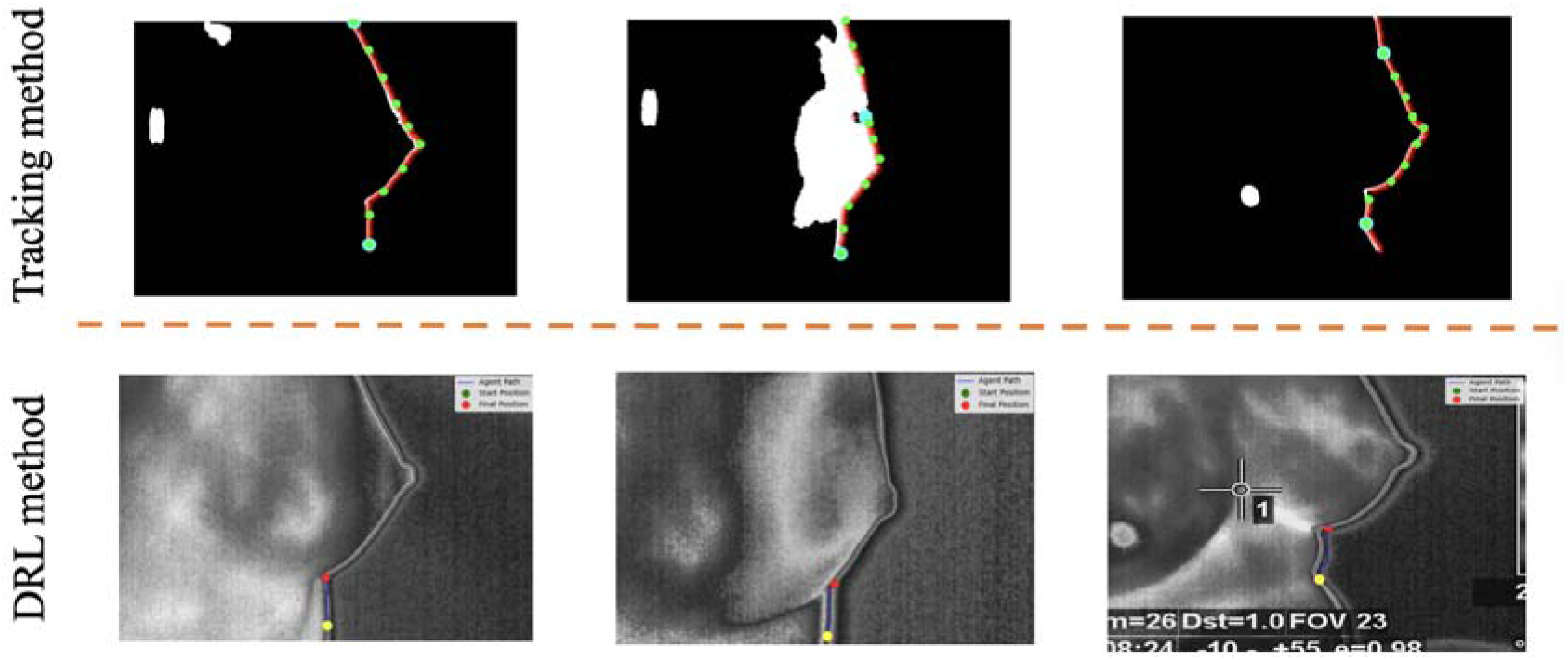
Salient point correction by Deep RL. The cyan dot (top row) indicates the salient point identified through the tracking method. The red point (bottom row) shows the corrected location by Deep RL.

Fig. 10.1 illustrates typical detected anterior (external) boundaries in lateral views. The initial landmarks are in red, and the black curves are the detected breast border. The images on the top row (complete shapes) belong to cases where the whole breast is visible from the lateral view. In the bottom row, however, part of the tissue is behind a barrier (incomplete shapes).

**Figure. 10.**
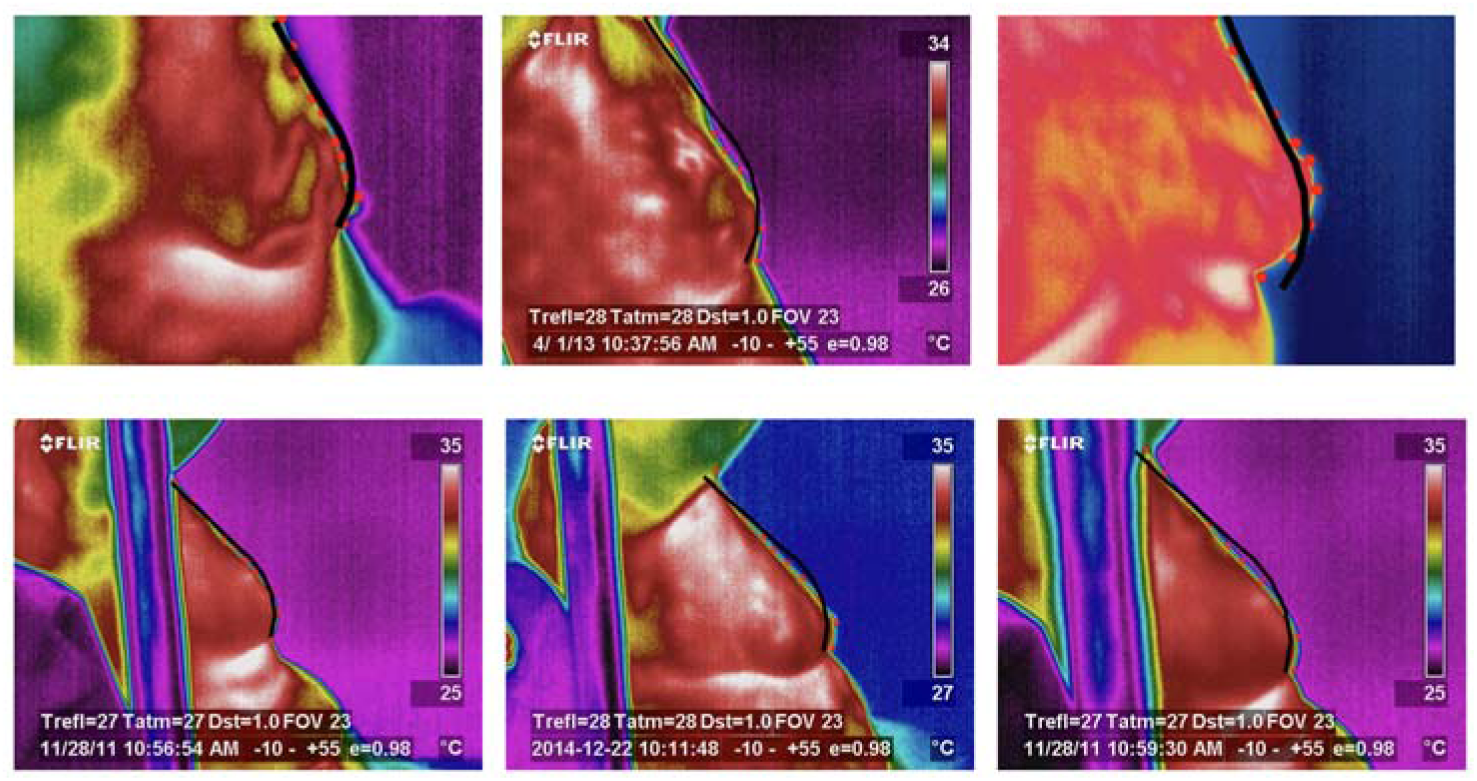
Typical results of the boundaries (black curves) of the breast of complete (top row) and incomplete (bottom row) shapes.

Boundary delineation continues with the definition of the posterior (internal) boundary modeled by a semicircle. The complete contour defines a binary mask of the breast region that is applied to the original thermal image to isolate the breast tissue. The segmentation remains part of the image where important thermal characteristics of the tissue exist. This multi-stage segmentation demonstrates the robustness of the approach across varying breast morphologies and thermal patterns (Figure 11).

**Figure. 11.**
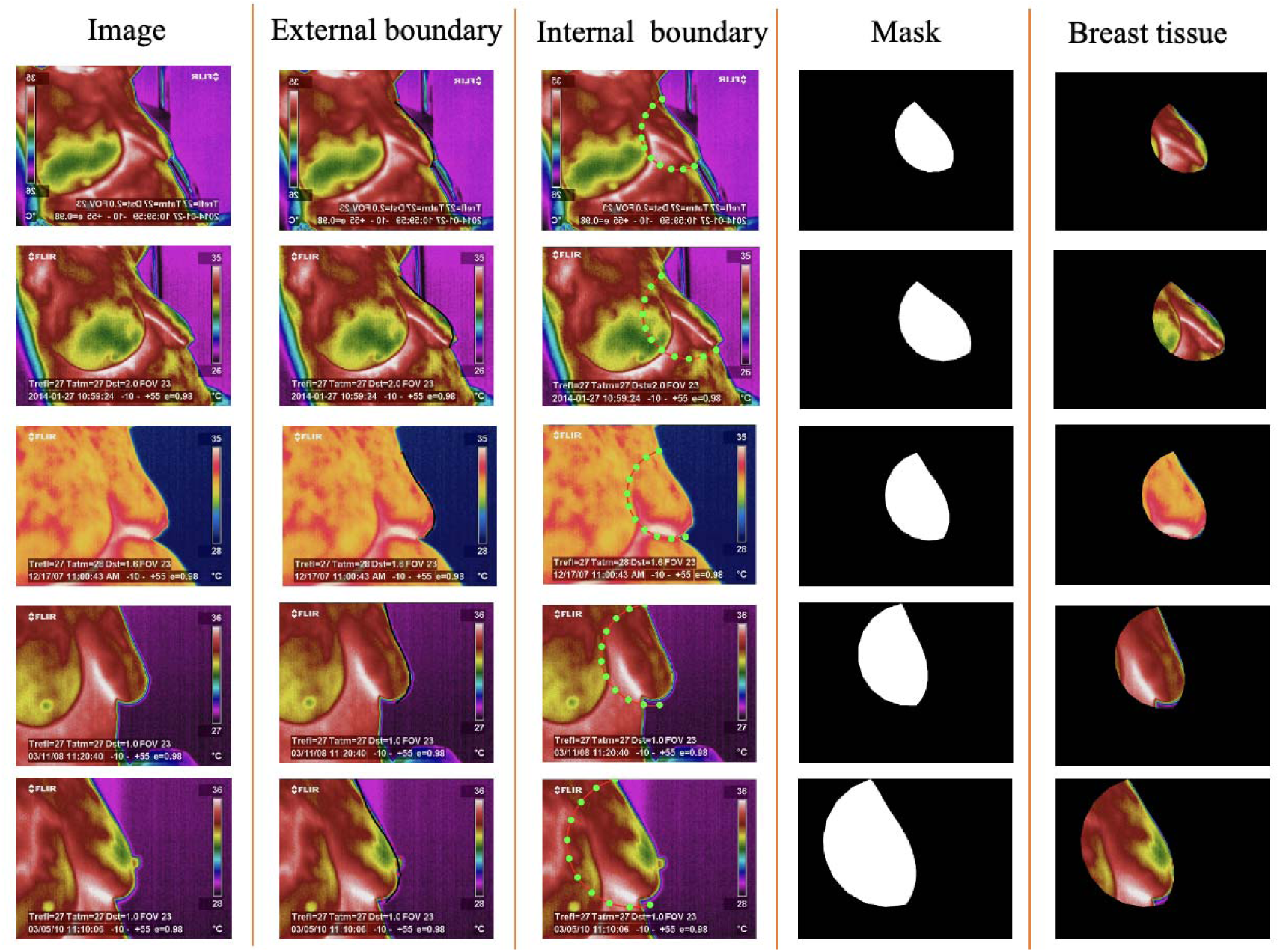
Typical results of breast region segmentation with varying breast morphologies and thermal patterns. Columns from left to right: original thermal images, detected external (anterior) boundaries, internal (posterior) boundary detection, breast binary mask, and segmented breast tissue.

To analyze the impact of the ROI on classification performance, we prepared representative examples from image categories used in our classification experiments (Fig. 12). The first row displays the original thermographic breast images with a complete field of view used in our initial classification scenario. The second row shows manual definitions of bounding boxes around the tissue regions. The third row demonstrates the results of our segmentation method, where breast tissue boundaries are precisely delineated. The following section discusse the quantitative impact of these different preparation methods on classification performance.

**Figure. 12.**
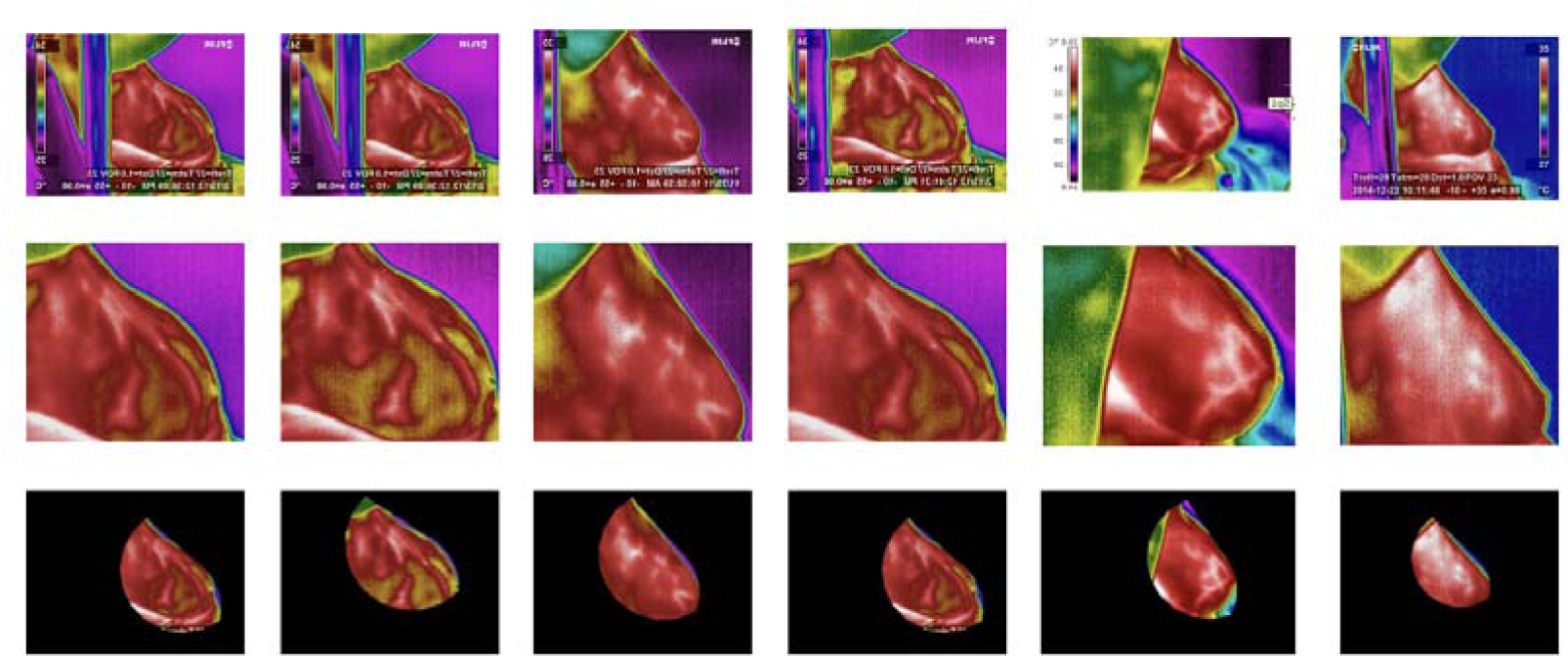
Different schemes to define the ROI of the breast. The first row displays original thermographic breast images. The second row is an ROI around the breast defined manually. The third row is the outcome of our method.

The proposed segmentation method has a clear advantage over conventional techniques, such as Active Contour and Fuzzy C-Means. While the Active Contour method faces challenges, including low-gradient boundaries and noise in lateral thermal images (Fig. 13a), the Fuzzy C-Means method (Fig. 13b) achieves acceptable segmentation results. Since these methods depend on manual interaction, they are not scalable. However, our method is a fully automated and robust solution. Therefore, our approach is well-suited for large-scale applications, including automated breast tissue analysis systems.

**Figure. 13.**
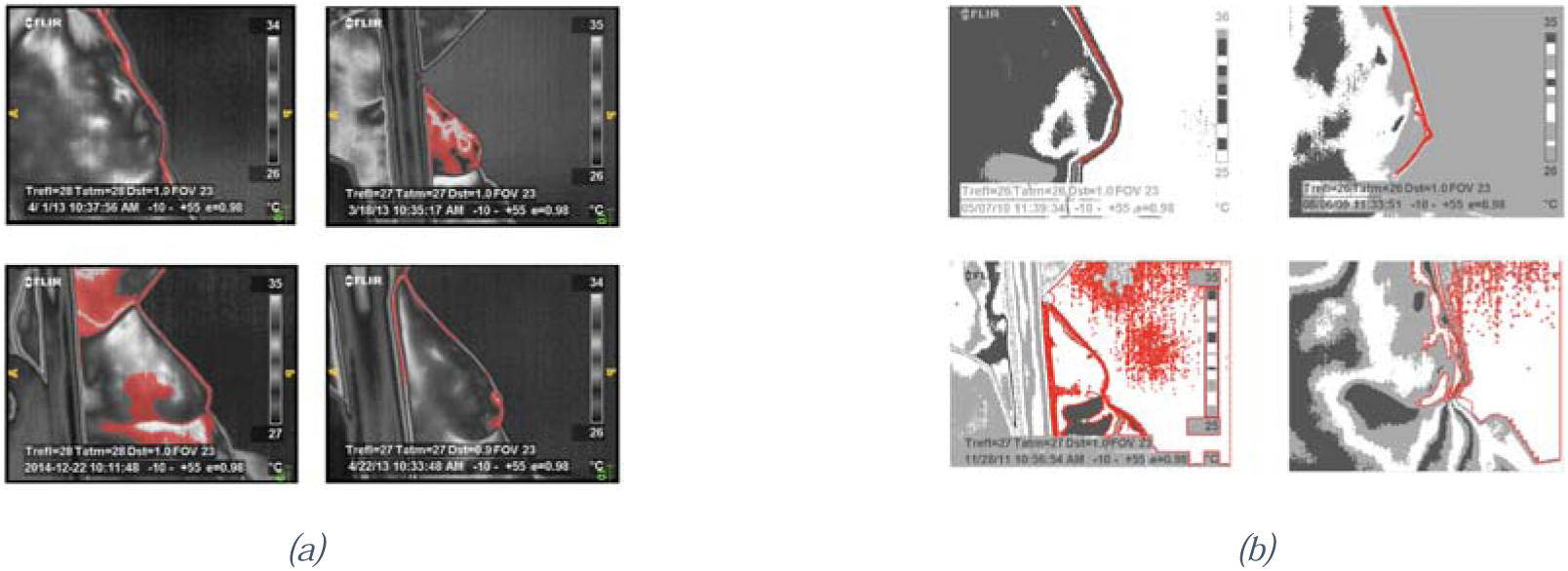
Typical breast segmentation using (a) geometric active contour and (b) fuzzy C-Means. The cases include complete and incomplete shapes with various scaling.

### 4.3 The RL performance

We used performance metrics in the learning process of the RL to confront th fluctuating nature of reinforcement learning (RL) losses. We tracked three key metrics: (1)

Euclidean gaps between predicted and true landmarks, (2) step counts to hit targets, and (3) total rewards per episode. Together, they reveal how robustly our RL agent learns and evolves.

Figure 14 shows the performance of the training of the RL agent. The three key metric are: steps taken per episode (yellow line), initial distance from the target landmark (green line), and final distance after correction (blue line). The curves in Fig. 14 show consistent improvement throughout training. Figure 14 reveals that the agent’s training is practical, since the number of steps required to reach the target decreases. This trend enables the agent to refine its correction strategies progressively through continuous learning.

**Figure. 14.**
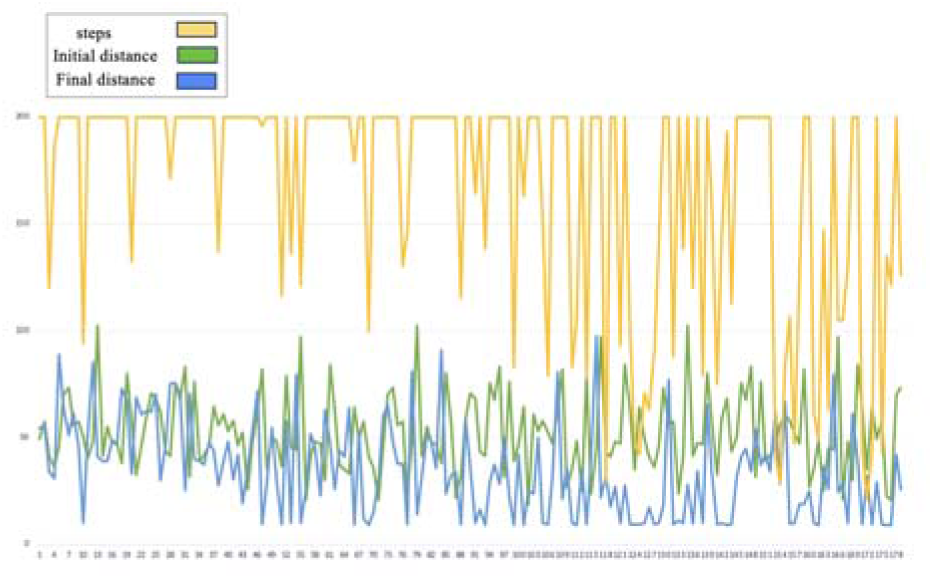
The performance of the RL agent over several episodes. The yellow line represents the number of steps taken per episode with a decreasing trend that suggests a more efficient pathfinding. The green line is the initial distance from the target landmark. The blue line is the final distance achieved after correction.

Our agent was successful in adjusting landmarks toward targets, achieving a remarkably low average distance error of 3.49 pixels. This consistent precision, even in complex cases, confirms its efficacy in minimizing localization errors.

Figure 15 maps learning progress via cumulative rewards. Initial episodes show deep negative rewards (down to -12,000), indicating poor early decisions. Post-episode 50, rewards stabilize and trend upward as performance improves. The diminishing amplitude of oscillations over time indicates that the consistency in landmark correction strategies also improves. There is a gradual increase from deep negative rewards to zero that reveals the efficiency of the method.

**Figure 15.**
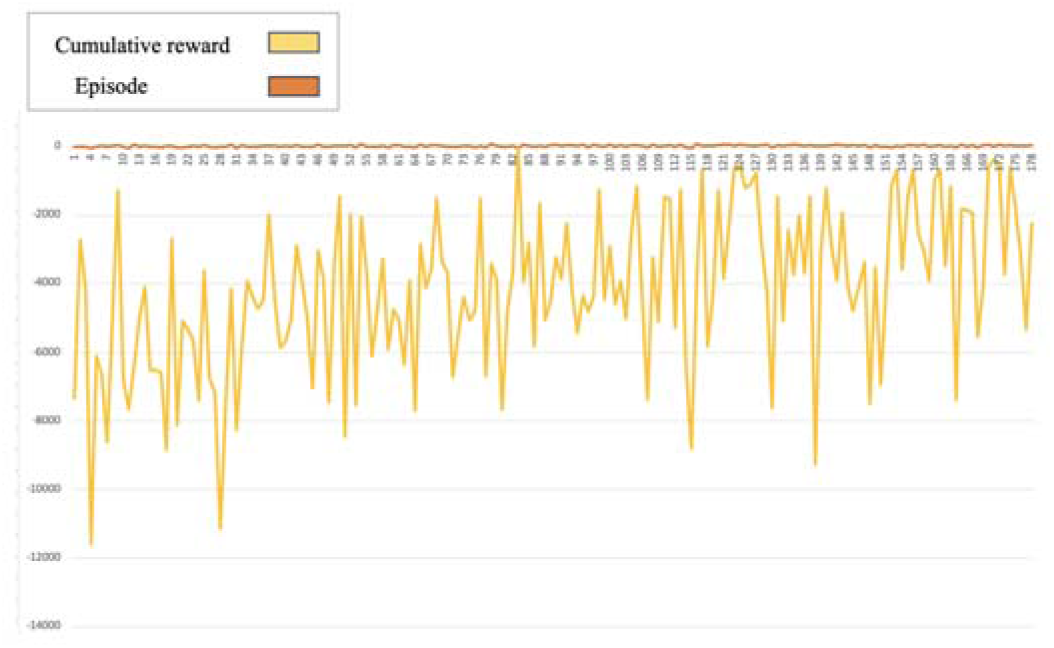
Learning progression of the reinforcement learning agent over training episodes. The yellow line represents the cumulative reward per episode, while the orange line indicates episode numbers.

### 4.4 Classification results

In the classification phase, we used an SVM with a linear kernel with features from the segmented region. The results of the classification further uncovered that accurate tissu segmentation plays a vital role in classification accuracy. Using basic GLCM features, th classification results in three regions were compared: (1) the whole thermography image, (2) a cropped ROI defined around the breast tissue manually, and (3) the ROI obtained by the proposed segmentation method. Our method outperformed both original and cropped images. While classification using the original images achieved an accuracy of 58.9%, using the cropped images dropped to 53.4%, and our approach achieved a remarkable 74.14% accuracy (Fig. 16). The improvement in classification accuracy demonstrates the impact of defining the tissue’s ROI on classification performance.

**Figure 16.**
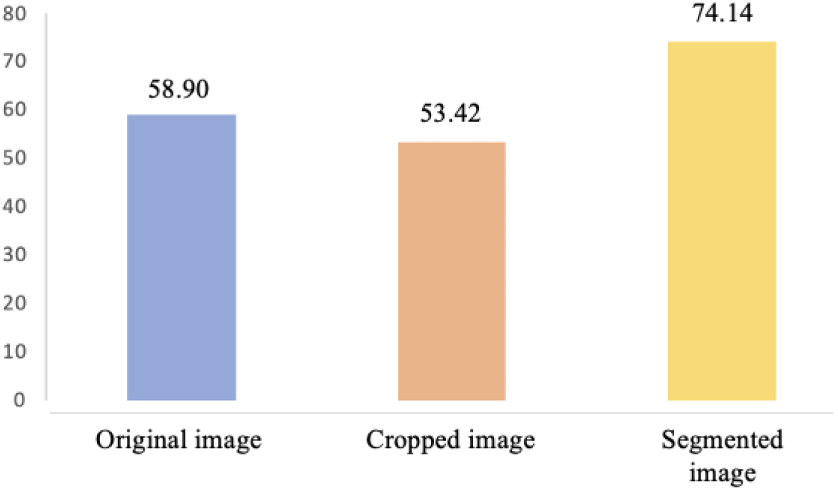
Classification accuracy of the SVM classifier using features from three different regions.

### 4.5 Comparison with recent research

Our approach shifts from frontal to multi-angle lateral thermography (30°, 45°, 90°), capturing richer anatomical data despite slightly lower accuracy (Table 1).

**Table 1:** Comparison with the proposed method and previous studies.

| Aspect | Previous Studies | Our Approach |
| --- | --- | --- |
| **Image views** | Frontal thermography | Lateral thermography<br>(30°, 45°, 90°) |
| **Machine Learning**<br>**Methods** | SVM (RBF and gaussian<br>kernel) [12,13,15]<br>Decision Trees [15]<br>K-star [14] | SVM (Linear kernel) |
| **Feature Extraction** | Wavelets [12]<br>LBP and HOG [13]<br>Gabor [16] | GLCM |
| **Deep Learning**<br>**Methods** | CNN [17]<br>Custom CNN [ 18]<br>FastLeakyResNet-CIR [19]<br>Inception MV4 [20] | ----- |
| **Reported Accuracy** | 77.8% to 100% | 74.14% |
| **Key Differentiators** | Frontal view<br>Dynamic and static data [14]<br>Hybrid models [19,20] | Lateral view<br>Multi- angle data<br>Simplified model |

## 5. Discussion

This study focused on classifying thermal images with lateral angles of 35°, 45°, and 90°. Most previous studies processed frontal views; however, lateral views are more complex due to existing occlusions and variable tissue scales (Fig. 17). Our method is scalable in addressing these challenges, making it beneficial for applications such as remote monitoring.

**Figure 17.**
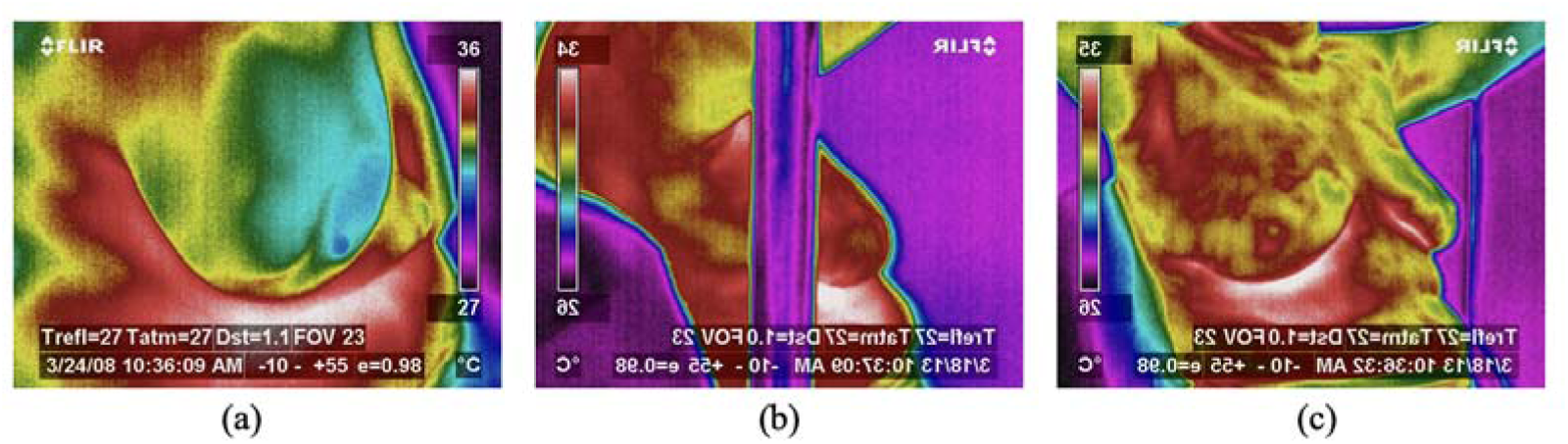
Typical challenges in lateral view data. (a) Zoomed-in image, (b) an image with an obstacle, and (c) a zoomed-out image by focusing on the internal lateral view of the right breast.

Our approach leverages and lateral thermography (LIME/LIMD configurations). Unlike frontal or views, these angles penetrate hidden internal breast regions, delivering superior tissue clarity. Consequently, we gain enhanced visibility into deep tissue architecture— extracting diagnostic data inaccessible through single-perspective imaging. A new viewing angle contributes unique and complementary insights into the diagnostic process.

Our method has the advantage of achieving robust segmentation and landmark detection with relatively small datasets. Unlike traditional deep learning approaches, which rely on large-scale annotated datasets, the reinforcement learning framework and statistical shape modeling used in this study demonstrated effective performance with just 14 training images initially and 77 in the expanded dataset. This approach is data-efficient in medical image processing when resources are limited. Dependence on large datasets often restricts the scalability and practicality of a method.

Our reinforcement learning approach shows real potential for landmark detection. In early tests with just 24 images (14 for training, 10 for testing), it performed well, even when landmarks started within 60 pixels of the targets, a challenging scenario. The RL agent trained on 128×128 crops successfully learned to reach target landmarks in 30,000 episodes. Figure 18 tracks how distance errors decrease progressively during training. This convergence proves that the learning stage is successful and robust even in challenging scenarios. Moreover, the final distances range from 5 to 7 pixels.

**Figure 18.**
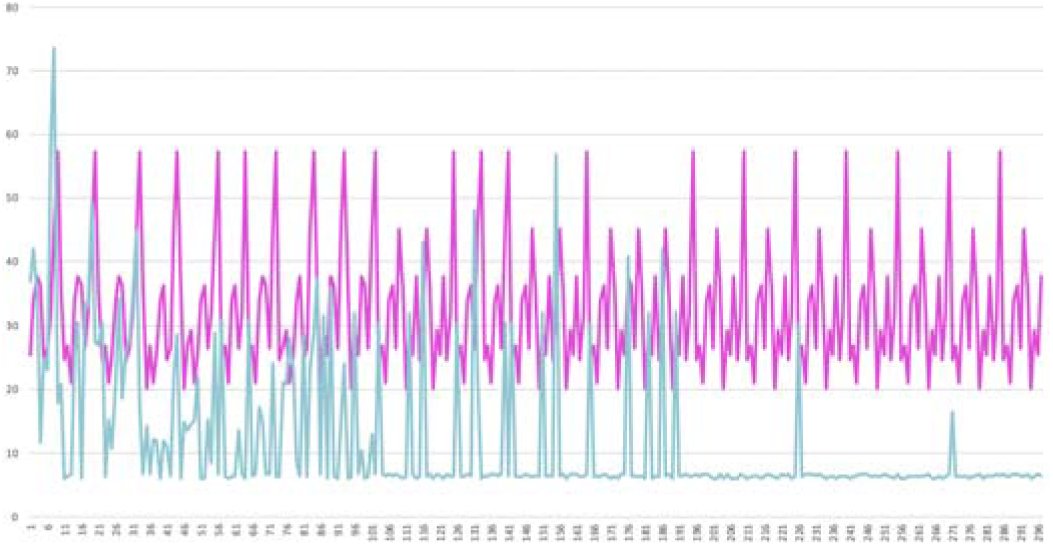
Distance convergence analysis during training of reinforcement learning. The magenta line represents the initial distances between the starting and target points, ranging from 20 to 57 pixels, while the blue line shows the final distances achieved after agent correction.

In a limited dataset with sufficient episodes and total steps, the reinforcement learning agent demonstrated strong performance in landmark detection across our test dataset. The agent successfully learned to navigate from various starting positions to target landmarks (Fig. 19, first row), even in challenging scenarios where the initial position is in complex regions, such as the image legend area (Fig. 19, second row).

**Figure 19.**
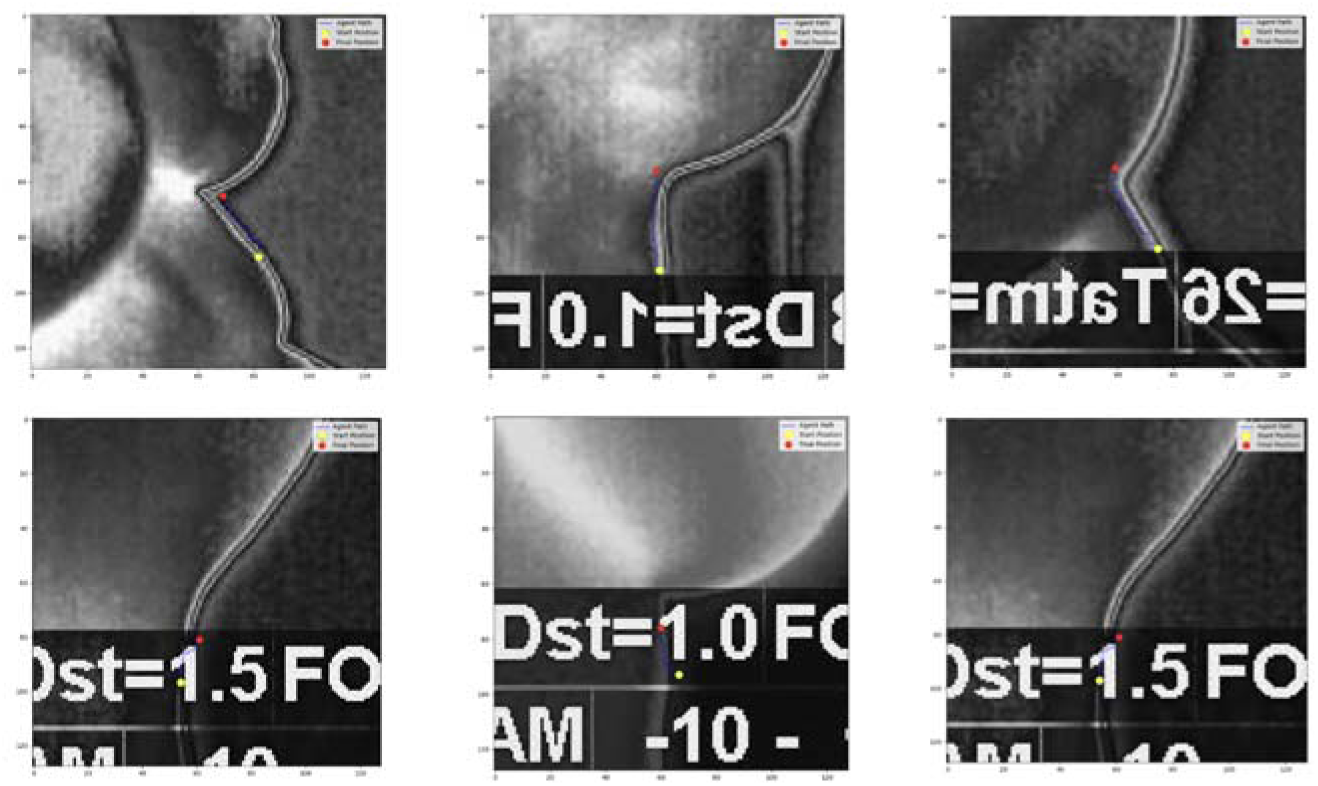
The landmark detection performance of reinforcement learning agent. The yellow point is the startin point, and the red point is the target point. (Top row) The agent successfully navigates from various initial positions to accurately identify target landmarks. (Bottom row) The agent demonstrates robust landmark detection even when starting from challengin locations, such as the image legend area.

In another experiment, we increased the training images to 77 and the initial distance to a maximum of 199 pixels to induce computational complexity. With only 30,000 training steps available, far fewer than ideal, our agent showed early learning promise but struggled to correct errors over large distances. Our solution is an adaptive training pipeline. We started with shorter 110-pixel gaps, then methodically ramped up step counts to bridge the capability gap. After optimization, the payoff was clear, especially in how steadily the agent nailed landmark correction despite tight resource limits.

The study also revealed two types of results from the DRL algorithm: landmarks successfully detected precisely at the target location (Fig. 20), and landmarks detected near the target point but requiring further refinement (Fig. 21). When initial detections landed near targets without perfect alignment, our Statistical Shape Model (SSM) fine-tuned their positions. This refinement boosted final accuracy significantly. The outcome confirms SSM’s critical supporting role in RL-based correction, proving multi-technique integration delivers truly robust solutions.

**Figure 20.**
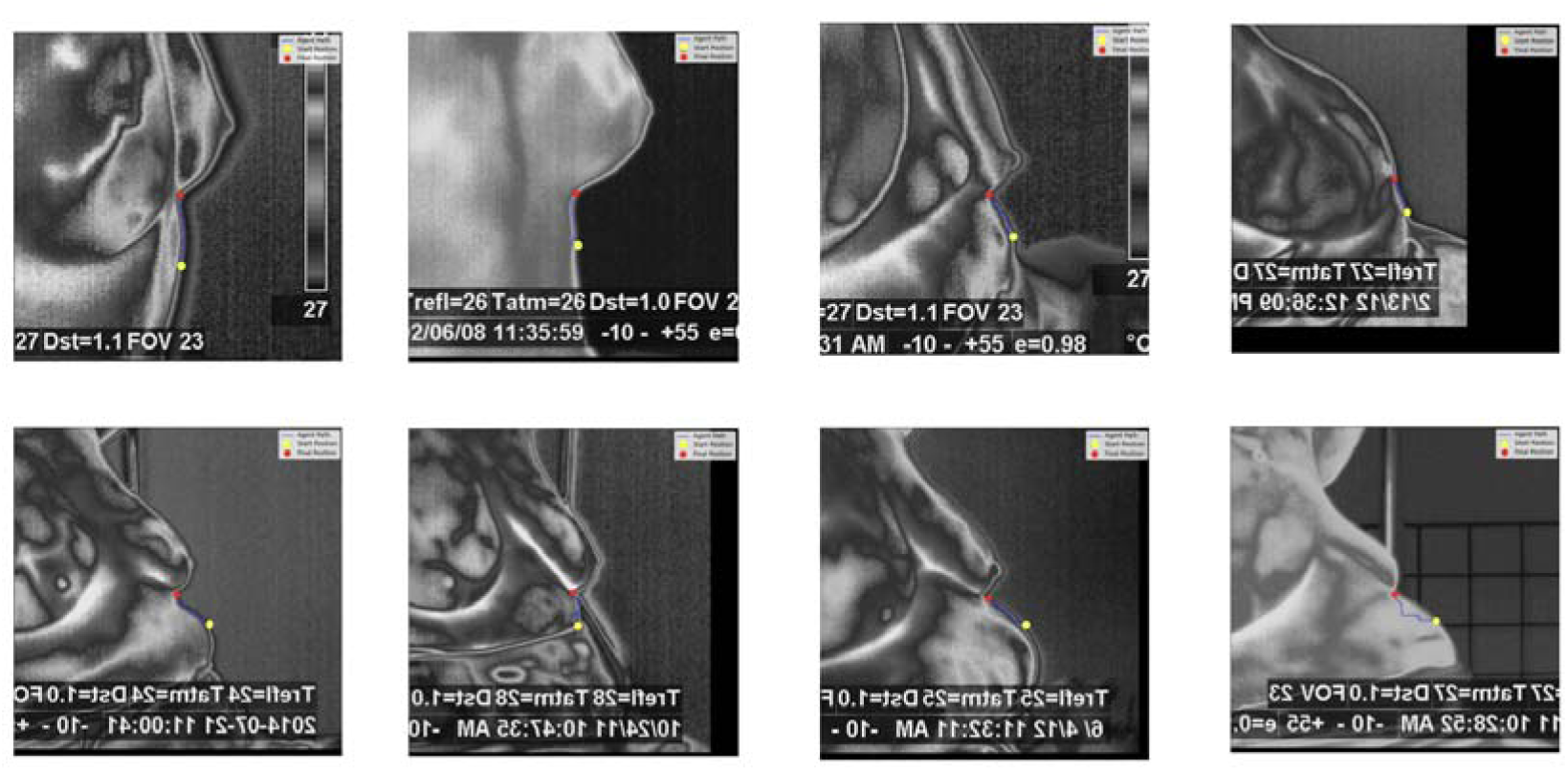
Results of the DRL algorithm successfully detecting the target points. The yellow point represents the starting position, and the red point indicates the location reached by the DRL algorithm.

**Figure 21.**
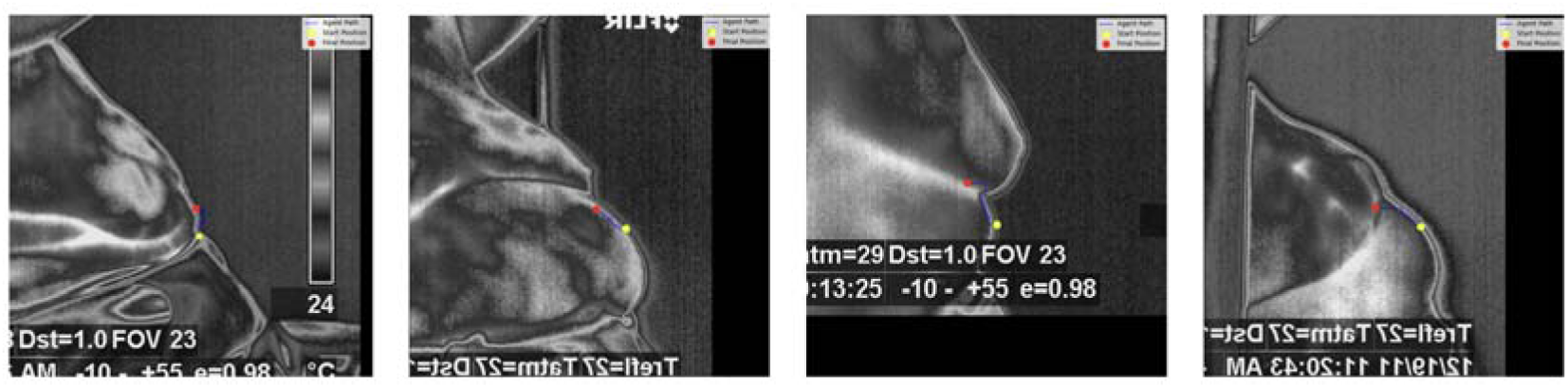
Results of the DRL algorithm where the target points were not detected precisely, but the detected points are close to the target locations. The yellow points is the starting position, and the red point is the location reached by the DR algorithm.

Our DRL results look promising, but we spotted three key constraints: training data volume matters, landmark starting-to-target distance plays a role, and search area size impacts outcomes. Real-world use might demand serious computing power for lengthy training runs, or clever tweaks to handle bigger distances and search zones. While the DRL method shows great potential, further improvements in training efficiency and resource utilization are necessary for widespread clinical adoption.

Beyond the technical advances, our work highlights real clinical value. Accurately segmenting lateral thermal images could enable new remote breast monitoring tools and diagnostic AI systems. Capturing and segmenting internal breast structures from side angles is a genuine leap forward for thermography. Next steps are to streamline RL training, add more diverse patient data, and combine other imaging techniques to boost diagnosis precision.

## 6. Conclusion and Future Works

Our framework advances breast cancer detection in lateral thermography by solving view-specific problems. Integrating preprocessing, scanning, tracking, RL, and SSM enabled reliable automatic segmentation and classification. Our RL landmark correction delivered significant gains in breast boundary precision—especially for tricky cases. Boundary-based segmentation became the unsung hero too, driving classification accuracy to 74.1% versus 58.9% for original images and 53.4% for cropped versions. This result proves two truths: pixel-perfect segmentation is non-negotiable for trustworthy diagnosis, and ML’s integration marks a potential turning point for medical imaging.

Moving forward, we are strengthening accuracy, automation, and clinical deployment readiness in our framework. First, we are creating a multi-agent DRL to autonomously identify all boundary points—removing intermediary steps while improving precision. Second, we are broadening datasets to encompass greater diversity across patient demographics, imaging angles, and tissue properties.

We are deploying CNNs and transformers to upgrade classification through end-to-end feature learning. These architectures promise more robust feature representations. Independently, we will extend the framework to create 3D breast reconstructions from multi-angle thermal images.

These 3D rebuilds give doctors richer anatomical insights, enhancing both diagnostic precision and visualization. Speed matters too because real-time results make our algorithm suitable for clinical use. To get there, we will refine algorithms, accelerate hardware, and deploy resources smarter.

Finally, we are giving landmark placement a more brilliant start, crucial for handling tricky cases like hidden tissues, blurry shapes, or low-quality scans. This upgrade will make-or-break reliability across all imaging conditions. Together, these steps are building a flexible, scalable system that could reshape how clinics use thermography to detect breast cancer.

